# Beyond Better Models: Barriers and a Path Forward in Generalisable TCR–Antigen Specificity Prediction

**DOI:** 10.64898/2026.10.12.758456

**Authors:** Ilija Ilievski, Agne Antanaviciute, Graham Ogg, Hashem Koohy

**Affiliations:** Medical Research Council Translational Immune Discovery Unit (MRC TIDU), Weatherall Institute of Molecular Medicine, University of Oxford, United Kingdom; MRC WIMM Centre for Computational Biology, Weatherall Institute of Molecular Medicine, Oxford, United Kingdom

## Abstract

T cells are at the centre of adaptive immunity, protecting the organism against infection and cancer. T-cell receptors (TCRs), expressed on the surface of T cells, enable the recognition of target cells through interactions with peptides presented by major histocompatibility complex (pMHC) molecules. Reliable, large-scale mapping of TCRs to their cognate pMHCs remains a holy grail of modern systems immunology, with far-reaching implications for cancer immunotherapy, vaccine development, and the understanding and treatment of autoimmune diseases. While exhaustive experimental mapping remains intractable due to the enormous combinatorial sequence space of TCRs and pMHCs, computational approaches offer a promising alternative. Consequently, we have witnessed increasingly sophisticated computational models of TCR–antigen specificity, from small neural networks to deep generative models and large protein language models. However, several independent benchmarks have shown that these models, despite their size and sophistication, largely fail to generalise to TCRs or pMHCs not present in the data available during model development. Here, we investigate possible reasons for this and suggest a path forward to generalisable TCR–pMHC prediction. First, our analysis of the largest public repositories of TCR–pMHC data reveals significant biases towards a small number of antigens and MHC molecules, sparse sampling of TCR sequence space per pMHC, as well as inconsistencies and inaccuracies in data collation. Second, we show the importance of carefully setting up model evaluation to measure discrimination based on antigen specificity instead of confounding factors present in the evaluation data. Finally, we highlight the need for and propose how best to match the evaluation metric with the specific downstream application of the predictions. Our investigation suggests that we, as a community, need more representative and better curated data, and a robust set of metrics and benchmarks to guide future model development towards generalisable TCR–pMHC specificity prediction.

## 1 Introduction

Predicting the antigen specificity of any T-cell receptor (TCR) has been a long-standing goal in adaptive immunology research. Achieving this goal may advance cancer immunotherapy [1], improve the diagnosis and treatment of autoimmune diseases [2–4], and accelerate vaccine development [5]. The most studied TCRs commonly recognise antigens in the form of peptide–MHC (pMHC) complexes, in which peptides are presented by major histocompatibility complex (MHC) molecules, known in humans as human leukocyte antigens (HLAs) [6]. Next-generation sequencing (NGS) of immune repertoires and high-throughput screening (HTS) have enabled the collection of TCR sequences and experimentally characterised TCR–pMHC interactions at an unprecedented scale [7, 8]. Yet, the enormous combinatorial diversity of an estimated 10^18^ possible TCRs [9, 10] and 10^14^ peptides [11], further compounded by more than 45,000 known HLA alleles [12], makes exhaustive experimental characterisation of TCR antigen specificity intractable. However, the growing availability of experimentally characterised TCR–pMHC data instead provides an opportunity to computationally infer TCR antigen specificity.

The opportunity has attracted considerable interest from the computational biology community, leading to the development of numerous models for predicting TCR specificity over the past decade [13–17]. However, several independent benchmarks have shown that these models struggle to generalise, with performance deteriorating when either the TCR or antigen is absent from the data used to develop the model [18–22]. This limited generalisability restricts their application for donor-derived TCRs and novel or underrepresented disease-relevant antigens, hindering their clinical translation.

In this work, we identify two key obstacles to generalisability, the quality and diversity of the available TCR–pMHC data and how models are evaluated. High-quality data comprising verified TCR–pMHC pairs with complete sequence information are essential, as the available data constrain whether a model learns transferable biological determinants of TCR–antigen specificity or data biases. While data limitations have received increasing attention [23–27], the impact of model evaluation in achieving generalisability remains underexplored. A robust evaluation setup that matches the biological question at hand is critical, as it guides model development and reveals whether reported performance is driven by antigen specificity or confounding factors in the evaluation data. Addressing these limitations in data and evaluation may therefore be as important as developing novel models.

We investigate these issues through three dedicated analyses. We first systematically analyse the composition, quality, and completeness of public TCR–pMHC data and highlight specific annotation and data ingestion problems (Section 2). We then show that the evaluation setup alone can radically alter the conclusions drawn about model performance (Section 3). Specifically, we examine whether shared specificity TCRs (with the same cognate antigen) are considered by representative TCR similarity methods to be more similar to each other than to TCRs used as negatives. Varying the composition of the shared specificity and negative TCRs groups shifts performance from perfect discrimination to approximately random or even strongly reversed ranking. Finally, we highlight how different evaluation metrics capture distinct aspects of model performance, including global ranking, retrieval of shared specificity TCRs within a limited screening budget, and the quality of antigen-specific TCR clustering (Section 4). Based on these analyses, we provide recommendations for data collection, benchmark construction, and metric selection that aim to better align model evaluation with the downstream application and guide future model development towards generalisable TCR–antigen specificity prediction.

## 2 Data Requirements for Generalisable TCR Specificity Prediction

All machine learning models fundamentally require data representative of the data space we want the models to generalize across [28]. The two largest actively maintained sources of public TCR– pMHC data are IEDB [29] and VDJdb [30]. IEDB is the older and substantially larger repository, established in 2004, with a primary focus on immune epitopes rather than TCR specificity. It contains 1.9 million epitope records, of which approximately 290K are TCR–pMHC human records. IEDB data is curated by a team of experts and uniquely includes data on non-binder TCR–pMHC pairs. Although very limited, these negative data are particularly valuable for model development and evaluation. VDJdb was established 13 years later with a sole focus on collating TCR specificity data. Two thirds of VDJdb TCR–pMHC pairs and one third of reference publications are not present in IEDB [31]. VDJdb data are curated by a small team, and the curation process thus relies more heavily on data verification through computational pipelines [32]. VDJdb uniquely includes confidence scores, derived from heuristic rules, that aim to quantify the strength of evidence for each TCR–pMHC record. While recognising their foundational importance to the field, our quantitative and qualitative analysis of these two repositories reveals considerable data biases and sparsity that limit their representative coverage of the TCR–pMHC sequence space of interest. In this section, we demonstrate these limitations in IEDB data for human αβ TCR–pMHC interactions, alongside issues related to data annotation and integration. A corresponding analysis of VDJdb data is provided in Supplementary Figure S2.

First, our analysis revealed pronounced biases in both antigen and HLA representation. Antigen representation in IEDB is heavily skewed towards a small number of viral organisms (Figure 1A). In the dataset, 89.1% of all antigen records are from just four viral source organisms: Severe Acute Respiratory Syndrome Coronavirus 2 (SARS-CoV-2; 54.8%), Epstein–Barr Virus (EBV; 18.3%), Human Cytomegalovirus (CMV; 9.3%), and Influenza A virus (6.8%). Other clinically important non-viral targets, such as tumour-associated neoantigens and autoimmune-relevant antigens, constitute only a minor fraction of the remaining records. HLA representation is similarly skewed towards a small number of HLA alleles (Figure 1B) relative to their frequencies in the population [33]. Four HLA categories account for 89.4% of all records: HLA-A*02:01 (32.6%), broad HLA Class I (30.0%), broad HLA Class II (21.0%), and HLA-A*03:01 (5.9%). The HLA analysis reveals another major problem with the data, where more than half of the records are annotated only at the level of HLA class, while other records are annotated at the level of allele. Reconciling data with different levels of HLA resolution requires carefully implemented data preprocessing pipelines and may result in further loss of data.

**Figure 1.**
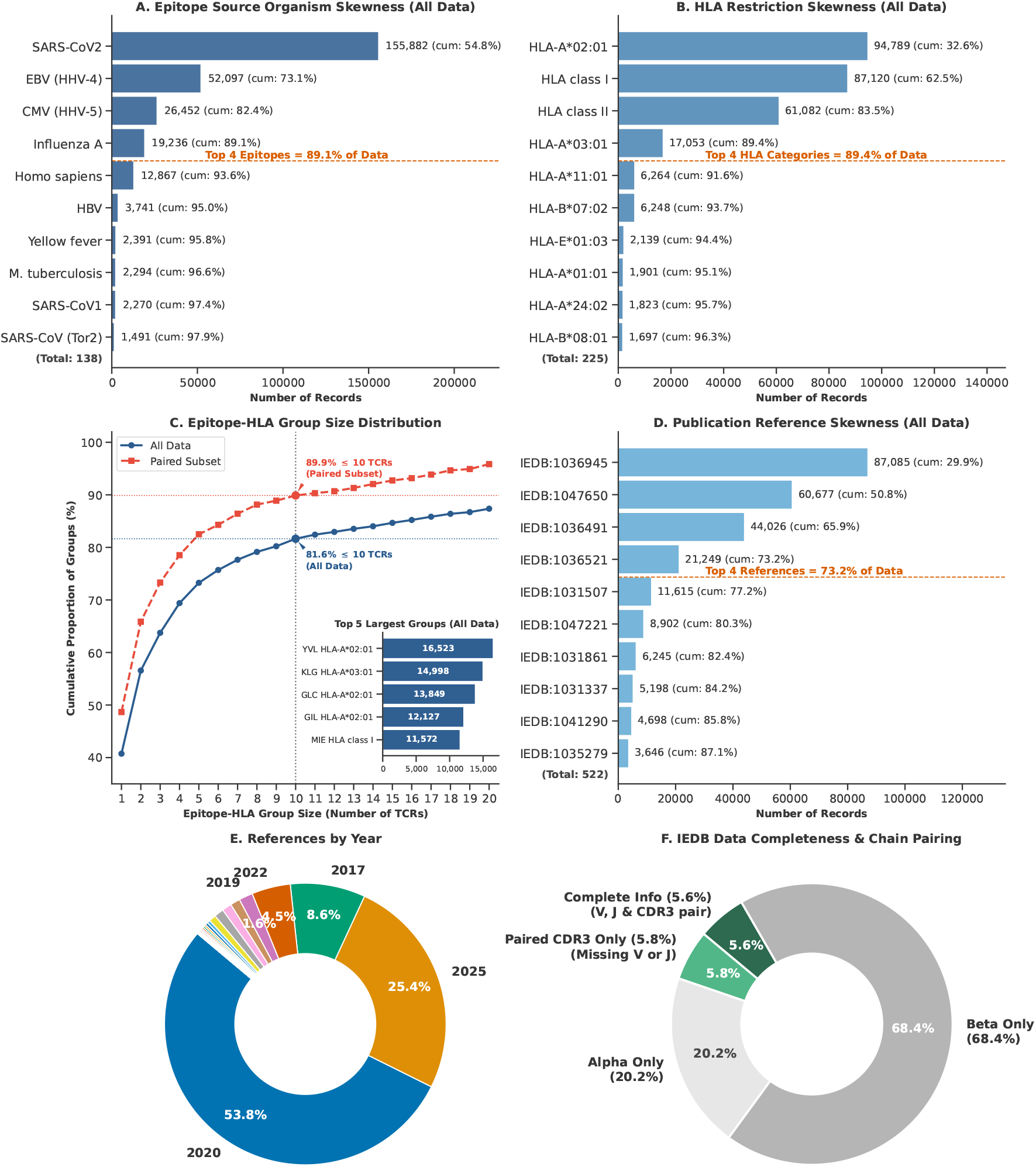
Quantitative analysis of data biases and annotation incompleteness in human αβ TCR–pMHC data from IEDB (≈290K records) (**A**) Epitope skewness. (**B**) HLA restriction skewness. (**C**) Cumulative distribution of number of unique TCR sequences per epitope–HLA pair. (**D**) Publication skewness. (**E**) Publication year distribution. (**F**) Data completeness and chain pairing.

Next, grouping unique TCRs reacting to each epitope–HLA pair in IEDB, including through binding, activation, or other unspecified interaction, revealed substantial differences in the number of TCR sequences per epitope–HLA pair (Figure 1C). Overall, 40.7% of epitope–HLA pairs are paired with only a single TCR record, and 81.6% with 10 or fewer TCRs. Considering only paired-chain TCRs, this sparsity is even more pronounced, with 89.9% of epitope–HLA groups containing 10 or fewer TCR sequences. At the opposite extreme, the five largest specificity groups each contain approximately 11,500–16,500 TCRs. Models therefore see many examples for a very small number of specificities, while receiving little information for most other specificities. This imbalance may bias models towards performing well on the few pMHC specificities that make up the majority of the data, while for sparsely represented pMHC groups there may simply be too few TCRs to learn their specificity sequence motifs [26].

Furthermore, the available TCR–pMHC data come from a small number of publications (Figure 1D). Just four studies contribute 73.2% of all IEDB records, with the two largest accounting for more than half of the entire dataset. This concentration means that many TCR records are not completely independent observations, but share the same experimental conditions, including assay platforms, donor cohorts, peptide libraries, sequencing protocols, and data processing pipelines. The publication bias is also reflected in when the data were published, with more than half of all records (53.8%) originating from studies published in 2020 and a further 25.4% from studies published in 2025 (Figure 1E). Another source of sampling bias is the anatomical origin of the TCRs, which may be collected from peripheral blood, tumours, or secondary lymphoid organs [23]. TCR repertoires can differ substantially between these compartments, with TCRs from tissue and lymphoid organs predominantly sampled from donors with disease. These differences may introduce additional confounding factors in the data. Models trained on these data may therefore learn study specific patterns that are shared across many records rather than genuine biological patterns of TCR–pMHC recognition. This may be difficult to identify and may result in inflated prediction performance when records from the same studies are present in both training and evaluation sets.

Finally, paired-chain sequence coverage in IEDB is also very limited (Figure 1F). TCR antigen recognition is determined by both the alpha and beta chains [27, 34], yet only 11.4% of all records contain paired CDR3-αand CDR3-βsequences. When requiring complete paired-chain information, including V and J gene annotations alongside the CDR3 sequences for both chains [35], the available data are further reduced to only 5.6% of all records. The overwhelming majority of records (88.6%) therefore contain only a single TCR chain, predominantly the beta chain (68.4%). This shows that only a small fraction of the almost 290,000 IEDB records is available for models to effectively learn the contribution of both chains to antigen specificity [36], including their interactions with each other and with the pMHC complex. Taken together, this analysis shows that the large number of records in IEDB overstates the effective coverage of the data space over which models are expected to generalise.

On the other hand, a TCR–pMHC record in a public repository does not necessarily represent definitive evidence of antigen specificity. VDJdb recognises this through its confidence score, which aims to quantify the strength of evidence supporting individual TCR–pMHC records. However, despite their strong correlation with functional validation [37], confidence scores are currently available for only approximately 8% of VDJdb records. The importance of distinguishing the strength of evidence is highlighted by the independent functional validation of VDJdb TCR–pMHC pairs by Messemaker et al. [37]. For two representative epitopes, Epstein–Barr virus GLC and SARS–Cov–2 YLQ, only approximately 30% of the evaluated receptors for GLC and 60% for YLQ showed the reported antigen reactivity upon reconstruction and functional testing. This discrepancy can partly arise because different experiments measure different aspects of TCR antigen reactivity. For example, pMHC multimer staining measures physical receptor engagement under multivalent conditions, whereas functional assays measure downstream T-cell responses such as activation or killing [38, 39]. Consequently, TCR–pMHC records broadly labelled as antigen-reactive can represent different experimental endpoints and different strengths of evidence for antigen specificity.

Experimental TCR sequences can be incompletely recorded when converted into standard database fields. Many computational models use the reported V-gene annotation to reconstruct the germline-encoded CDR1 and CDR2 sequences of a TCR. However, experimentally tested TCRs can contain engineered mutations within the CDR1 and CDR2 regions that cannot be captured when the database supports only canonical V-gene annotations, as is the case with VDJdb [40]. A model reconstructing these receptors from the deposited V-gene annotations would therefore receive CDR1 or CDR2 sequences that differ from those of the experimentally tested receptor. Database ingestion can also introduce direct errors into otherwise correctly reported TCR sequences. For example, during the ingestion of the high-throughput TCR data from Gaglione et al., CDR3-β sequences were duplicated into the corresponding CDR3-α fields for multiple VDJdb records [41]. Thus, even when the experimental specificity measurement itself is accurate, database limitations and ingestion errors can result in a database record that does not faithfully represent the experimentally tested receptor.

At the same time, given the vast combinatorial space of possible TCR–pMHC interactions and the restricted specificity of individual TCRs, a randomly selected TCR is often assumed not to recognise a given pMHC. However, even if this assumption is generally correct, such negatives provide limited information for learning the sequence determinants of antigen specificity. To learn what determines antigen specificity, particularly informative negative examples are TCRs that are highly similar in sequence to pMHC-reactive TCRs but have been experimentally shown not to recognise the same pMHC. We demonstrate the importance of these negatives in the following section, where TCR similarity methods readily distinguish NY-ESO-1-reactive TCRs from unrelated presumed non-binders, but fail to distinguish them from experimentally confirmed non-binders generated within the same mutational scanning experiment. IEDB is currently the only major public TCR–pMHC repository that includes experimentally reported negative interactions, but it contains only approximately 1,000 such records. These negative records are also subject to the same antigen, HLA, and study sampling biases identified above for the positive IEDB data. Thus, the available negative data are too limited for meaningful model development, particularly for learning the subtle sequence differences that distinguish TCRs that recognise a given pMHC from closely related TCRs that do not.

Recent efforts by public repositories show important progress towards improving the quality and consistency of TCR–pMHC data [25]. For example, the recent revision of adaptive immune receptor data in IEDB explicitly acknowledged the need for consistent curation suitable for computational analyses and introduced improvements towards this goal [31]. Such efforts can address many standardisation and database representation issues, including some of those identified above. However, improved curation cannot correct biases already present in the experiments and publications from which these databases are assembled. To examine whether the biases identified in this Section (Figure 1) primarily reflect legacy data, we repeated the analysis using only records from studies published in 2021 or later (Supplementary Figure S1). The same antigen, HLA, pMHC sampling, and study biases remain evident in these recent data. The four most represented antigen source organisms accounted for 94.5% of all records and the four largest studies for 80.1%. Although almost all paired-chain TCR records here contain complete chain information, they remain scarce at only 11% of the records. A corresponding analysis of VDJdb shows similar antigen, HLA, and TCR sampling biases: the four most represented antigen source organisms and HLA categories account for 84.1% and 83.1% of records, respectively, while 84.3% of epitope–HLA groups contain 10 or fewer TCR sequences (Supplementary Figure S2). These results show that the sampling biases identified in IEDB are neither restricted to legacy data nor specific to a single repository.

An important question is therefore how to generate representative datasets without exhaustive experimental TCR–antigen screening. One approach is to sample a broad range of peptides and HLAs, with representative TCRs for each specificity. However, Delaunay et al. [42] estimate up to 100 million epitopes would be needed to cover the point mutation-derived variants of the neoantigen space as an example. An alternative is suggested by Gfeller et al. [26], who recommends screening fewer but clinically relevant pMHCs more extensively, sampling diverse TCR sequences with closely related experimentally verified binders and non-binders. Such data would provide more information about the sequence determinants of antigen specificity, although their applicability to unseen pMHCs may remain limited. Ultimately, the appropriate balance between breadth and depth will depend on the intended application and could be guided by measuring how model generalisation improves as additional pMHC specificities and TCR sequences are sampled.

Overcoming these remaining limitations therefore requires a wider community effort to generate data across more diverse antigens and HLAs, with sufficient TCR sampling and experimentally validated informative negatives. Generating such data requires experimental, reporting, and funding priorities that consider the needs of computational model development. Importantly, these data limitations affect not only our ability to train trustworthy models, but also our ability to fairly evaluate whether models have learned antigen specificity. Ultimately, progress towards generalisable TCR–antigen specificity prediction depends on the representativeness and informativeness of the data available for model development and evaluation. The critical question is therefore not how much data we need, but what data we need.

## 3 Evaluation Data for Measuring TCR Specificity Model Performance

TCR antigen specificity can be inferred either by directly predicting whether a TCR–pMHC pair would interact or from the sequence similarity between two TCRs. In many applications the antigen recognised by a TCR is unknown, but we still want to better understand the TCR data by identifying groups of TCRs likely to share antigen specificity. This makes TCR similarity methods particularly useful. Furthermore, direct TCR–pMHC prediction remains limited by the small number of pMHCs available for training [18, 43].

We therefore focus on TCR similarity methods, where TCRs recognising the same antigen, which we refer to as shared specificity TCRs, are expected to be more similar to each other than to other TCRs used as negatives. We evaluate how well TCR similarity models can discriminate shared specificity TCRs from negatives, with the complete mathematical formulation and evaluation setup provided in the Supplementary. We use SCEPTR [13] and TCRdist3 [44, 45] as representative TCR similarity methods, together with a simple CDR3 edit distance baseline that sums the edit distances between CDR3-α sequences and between CDR3-β sequences. We show that the measured performance heavily depends on the composition of the groups of shared specificity and negative TCRs, because differences unrelated to antigen specificity can also make these two groups easy to discriminate. For this, we designed three experiments to illustrate how measured performance may reflect antigen specificity but also other systematic differences between the TCRs.

We use the NY-ESO-1-specific (A0201 SLLMWITQC) 1G4 TCR phage display mutational library from Croce et al. [46] as a controlled experimental system for constructing the three evaluation setups. The library contains a large number of experimentally measured binders and non-binders generated through mutational scanning (MutScan) of a small region of the 1G4 TCR. Specifically, the TCRs share the same sequence scaffold, the part of the TCR sequence that remains unchanged, while five central residues of CDR3-β are replaced by variable sequences of up to nine amino acids. We re-processed the raw FASTQ data from Croce et al. to identify binder and non-binder TCRs, with the complete processing details and code provided in Supplementary Section S2. This provided us with large groups of shared specificity

We first consider a conventional evaluation setup, where all TCRs known to recognise a certain antigen, in this case NY-ESO-1, are used as the shared specificity group and TCRs with no known reactivity to that same antigen are used as negatives (Figure 2, Group 1). All three TCR similarity methods achieve perfect discrimination in this setup, with an AUC of 1.00. However, closer inspection of the 43,530 shared specificity TCRs reveals that all except 10 originate from the Croce et al. mutational scanning experiment. These 43,520 TCRs therefore share the same 1G4 sequence scaffold and differ primarily within the small CDR3-β region varied during the mutational scan. In contrast, the negatives are drawn broadly from VDJdb, with diverse TCR gene usage, resulting in profound sequence differences between the two groups. The perfect discrimination can therefore be achieved by recognising the shared 1G4 scaffold, without necessarily discriminating TCRs based on NY-ESO-1 binding. Importantly, this does not mean that TCRs with no known shared specificity are necessarily not useful as negatives, for many TCR–pMHC pairs these may be the only negatives practically available. Rather, the perfect AUC may not by itself establish antigen-specific discrimination when the shared specificity TCRs are highly homogeneous while the negative set is so diverse, making these negatives uninformative and “easy” to distinguish from the shared specificity TCRs.

**Figure 2.**
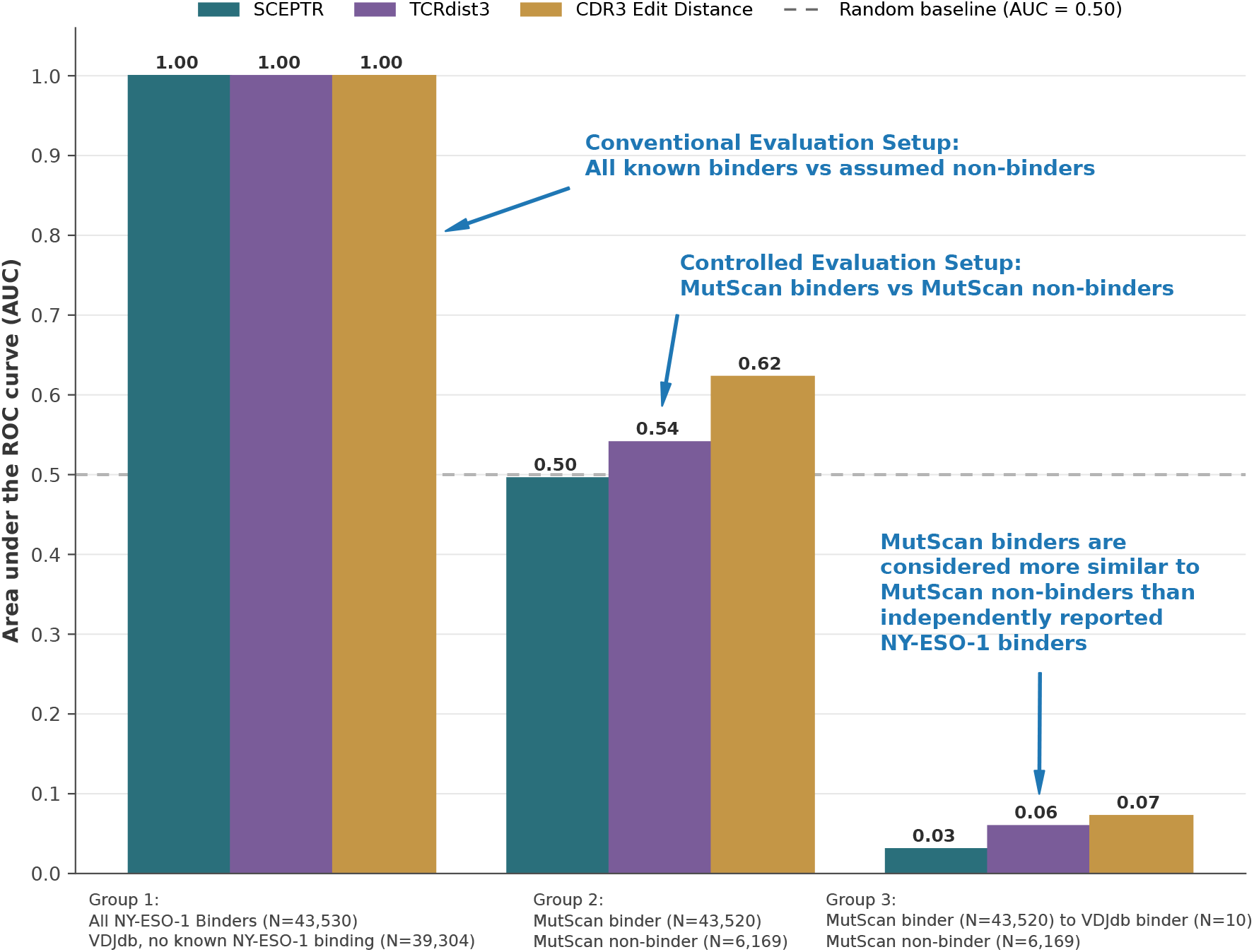
AUC performance of TCR similarity methods across three evaluation setups. **Group 1** compares all known NY-ESO-1 binders, including mutational scanning (MutScan) TCRs, with TCRs with no known NY-ESO-1 reactivity. **Group 2** compares MutScan binders with MutScan non-binders. **Group 3** compares distances between MutScan binders and other VDJdb NY-ESO-1 binders with distances between MutScan binders and MutScan non-binders. Bars show the AUC for SCEPTR, TCRdist3, and CDR3 edit distance (summed across both chains). The dashed horizontal line indicates random preformance (AUC = 0.50).

In the second evaluation setup, we restrict both groups to TCRs only from Croce et al. study (Figure 2, Group 2). The shared specificity group consists of the 43,520 experimentally verified NY-ESO-1 binders, while the negatives are the 6,169 experimentally verified non-binders. The two TCR groups therefore share the same 1G4 scaffold and experimental background, differing primarily in whether the TCR binds NY-ESO-1. This provides a more direct test of whether TCR similarity methods capture the specific residue differences associated with NY-ESO-1 binding rather than the shared sequence scaffold. Under this evaluation, the perfect discrimination observed in Group 1 disappears, with AUCs of 0.50 for SCEPTR, 0.54 for TCRdist3, and 0.62 for the CDR3 edit-distance baseline. Thus, when evaluated against closely related and experimentally verified non-binders, the three similarity methods discriminate poorly between NY-ESO-1 binders and non-binders.

The third evaluation setup aims to tests whether shared antigen specificity or sequence scaffold has a stronger influence on the measured TCR similarity (Figure 2, Group 3). Here, we compute the distances between the 43,520 MutScan binders and the 10 independently reported VDJdb NY-ESO-1 binders that were excluded from Group 2. We then compare these distances with those between the same MutScan binders and the 6,169 experimentally verified MutScan non-binders. This setup represents a practical specificity-discovery scenario, where the independently reported VDJdb binders are the TCRs with known NY-ESO-1 specificity, the MutScan non-binders serve as negatives, and we ask whether the MutScan binders, with their specificity treated as unknown, can be identified as NY-ESO-1 reactive. The independent binders therefore share NY-ESO-1 binding with the MutScan binders but not the 1G4 scaffold, whereas the MutScan non-binders share the 1G4 scaffold but not NY-ESO-1 binding. All three similarity methods produce strongly reversed rankings, with AUCs of 0.03 for SCEPTR, 0.06 for TCRdist3, and 0.07 for the CDR3 edit-distance baseline. This means that the experimentally verified NY-ESO-1 non-binders sharing the 1G4 scaffold are considered substantially more similar to the MutScan binders than genuine NY-ESO-1 binders from independent studies. This demonstrates that sequence scaffold similarity may dominate shared antigen specificity in these comparisons, explaining the perfect discrimination in Group 1, its disappearance in Group 2, and the reversed ranking in Group 3.

Although the phage display experiment provides a particularly clean example of sequence scaffold confounding antigen-specific discrimination, the same problem can arise from other systematic differences between shared specificity TCRs and negative TCRs [26]. The relevant confounding factors will depend on how the data for each antigen were generated and collected. For example, if most or all shared specificity TCRs for a particular antigen originate from a single donor, using TCRs from different donors as negatives may allow the two groups to be distinguished based on donor provenance rather than antigen specificity. In such cases, selecting the negatives from the same donor, where possible, would provide a more informative test of antigen-specific discrimination. In general, when constructing evaluation datasets, we should ask: what else, besides antigen specificity, systematically distinguishes these two groups? Any such differences provide alternative signals for discriminating between the groups, increasing the risk of inflated performance. Both TCR similarity models, and models that directly predict TCR–pMHC interactions are equally prone to this risk.

Near perfect performance for any antigen should therefore be interpreted with caution, particularly given the inherent biological and experimental noise in TCR specificity data. As demonstrated by Group 1, such performance can arise when systematic differences between shared specificity TCRs and negatives make the evaluation substantially easier than discriminating based on antigen specificity. This can be difficult to identify when model performance is reported only as an overall AUC averaged across multiple antigens [21]. A small number of antigens with perfect or near perfect discrimination can substantially increase the average AUC, while the potentially confounded evaluation responsible for these high values remains hidden within the aggregate score. We therefore recommend reporting and inspecting performance for individual antigens alongside the overall average, with particular attention to unexpectedly high performance and how the corresponding evaluation groups were constructed.

Fair model evaluation is particularly important because measured performance guides future model development by determining which approaches are prioritised for further improvement, making misleading performance particularly consequential. The three evaluation setups demonstrate that measured performance must be interpreted in the context of how the shared specificity TCRs and negatives were selected. A fair evaluation therefore requires negatives selected with consideration of other systematic differences between the two groups, so that successful discrimination provides evidence for antigen specificity rather than an easier alternative distinction. This requires understanding the provenance and composition of the evaluation data, rather than treating all TCRs with no known shared specificity as equally informative negatives. Ultimately, the biological question we want the model to answer should determine what data are used for evaluation.

## 4 Evaluation Metrics for Measuring TCR Specificity Model Performance

Selecting appropriate data is only one part of constructing an informative model evaluation. Different evaluation metrics measure different aspects of model performance, and the appropriate metric therefore depends on the biological or experimental question we want the predictions to answer. As model evaluation guides which approaches are retained and further improved, the choice of metric directly influences model development. For example, evaluating performance across the complete test set, identifying a limited number of candidates for experimental validation, and assessing the composition of a TCR repertoire represent distinct objectives that require different evaluation metrics. In the following, we examine commonly used evaluation metrics and discuss their appropriate application to these different objectives. We use TCR similarity methods as illustrative examples throughout this discussion, but the same considerations apply more broadly to other TCR specificity prediction methods.

The area under the receiver operating characteristic curve (AUC) is one of the most commonly used metrics for evaluating machine learning models and has consequently been widely adopted for evaluating TCR specificity prediction [18]. In the case of TCR similarity models, AUC measures how well the method ranks shared specificity TCRs ahead of the rest of the TCRs, termed negatives, across the complete test set. An AUC of 1 represents perfect ranking, 0.5 no better than random ranking, and 0 perfectly reversed ranking. AUC is therefore appropriate when the objective is to compare the overall discrimination performance of different models across the complete test set. However, global discrimination across the complete test set may not always correspond to the biological or experimental question of interest.

In many computational biology applications of TCR similarity models, we select a limited number of TCRs for further analysis or experimental validation based on their predicted shared specificity with a TCR of interest. In such applications, performance among the top *K* candidates is more relevant than discrimination across the complete test set. For these common applications, we propose reporting the enrichment factor at rank *K*, EF@*K*, which measures the proportion of shared specificity TCRs among the top *K* predictions relative to their proportion in the complete test set (Supplementary Section S3.3). An EF@*K* of 1 represents no better than random retrieval, while values above or below 1 indicate enrichment or depletion of shared specificity TCRs among the top *K*, respectively.

Figure 3 illustrates how AUC and EF@*K* can lead to different conclusions about model performance. Scenarios 1 and 2 have the same AUC of 0.75, but retrieve 80 and 50 shared specificity TCRs, respectively, among the top 100 candidates, corresponding to EF@100 values of 1.60 and 1.00. Conversely, Scenarios 3 and 4 have similar near-random AUCs of 0.48 and 0.51, while Scenario 3 retrieves 20% more shared specificity TCRs than expected by chance within the top 100 and Scenario 4 approximates random retrieval. AUC and EF@*K* therefore measure complementary aspects of model performance: AUC evaluates discrimination across the complete test set, whereas EF@*K* measures enrichment among the top *K* candidates.

**Figure 3.**
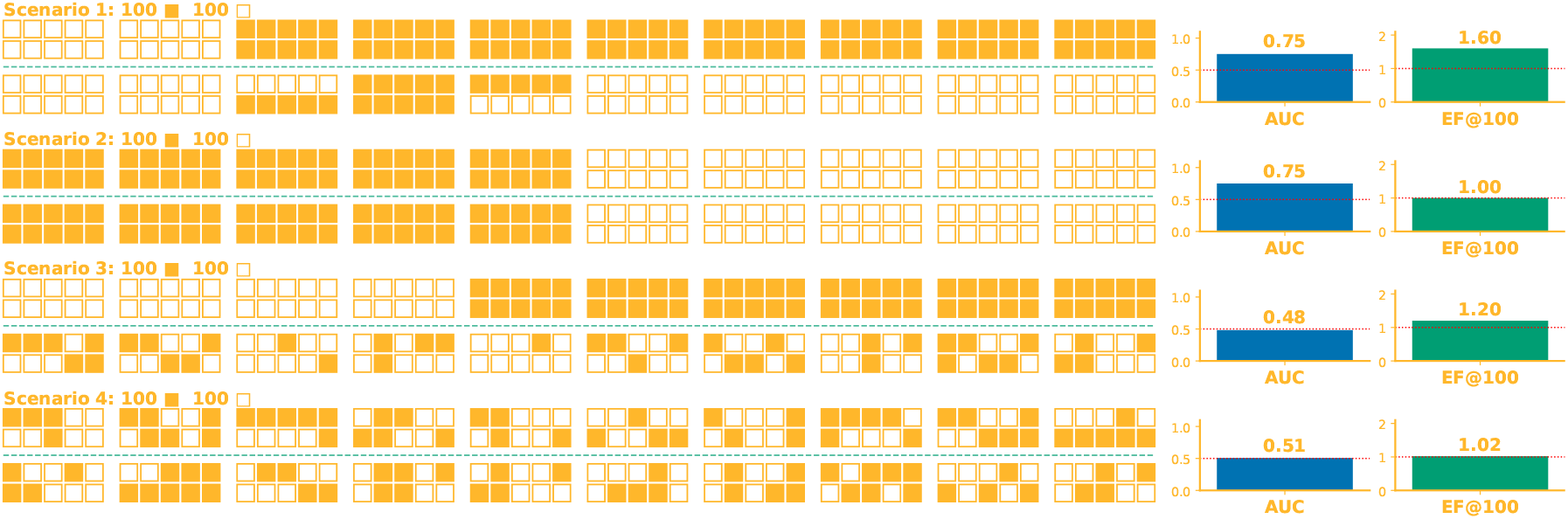
Choice of evaluation metric should reflect the intended experimental application. Four ranking scenarios compare the area under the receiver operating characteristic curve (AUC) with the enrichment factor among the top 100 predictions (EF@100). Each scenario considers, for a query TCR, 100 common-specificity TCRs (filled squares) and 100 negative TCRs (empty squares). The horizontal dashed line separates the top 100 ranked TCRs selected for experimental validation from the remaining 100 candidates.

Partial AUC (pAUC) provides an alternative when the objective is to evaluate performance within a restricted false positive rate (FPR) region [21]. By restricting the receiver operating characteristic curve to a predefined maximum FPR, pAUC scores the discrimination strictly at low FPR. However, a FPR threshold can be less intuitive to interpret in practical terms than a fixed rank *K*. For example, pAUC@0.10 evaluates performance while allowing up to 10% of all negatives to be retrieved, and therefore depends on the number of negatives in the test set, whereas EF@100 directly measures enrichment among the top 100 candidates. Standardised pAUC can further complicate this interpretation, as the transformation used to retain 0.5 as the random baseline compresses very poor performance towards 0.5. For instance, when pAUC is calculated up to a FPR of 0.10, a raw pAUC of zero is transformed to approximately 0.47 under the commonly used McClish standardisation [21, 47]. The standardisation and restriction of the pAUC to a fixed FPR also loses the interpretability of the score. The pAUC score no longer represents the probability of a shared specificity pair to be ranked higher than a negatives pair, as it does in the case of AUC. Additionally, pAUC can be unstable for TCR specificity groups containing only a small number of TCRs, as moving only one or two examples can substantially change the estimated area.

Figure 4 illustrates how these properties can lead pAUC and EF@*K* to give different views of model performance. Scenarios 1 and 2 have the same pAUC@0.10 of 0.74, but retrieve different numbers of shared specificity TCRs among the top 100 candidates, corresponding to EF@100 values of 4.80 and 3.00. Similarly, Scenarios 3–5 have pAUC@0.10 values close to 0.5, despite corresponding to substantial enrichment, complete depletion, and approximately random retrieval, respectively. Thus, pAUC may be appropriate when performance within a restricted FPR region is the question of interest, but its limitations should be taken into consideration.

**Figure 4.**
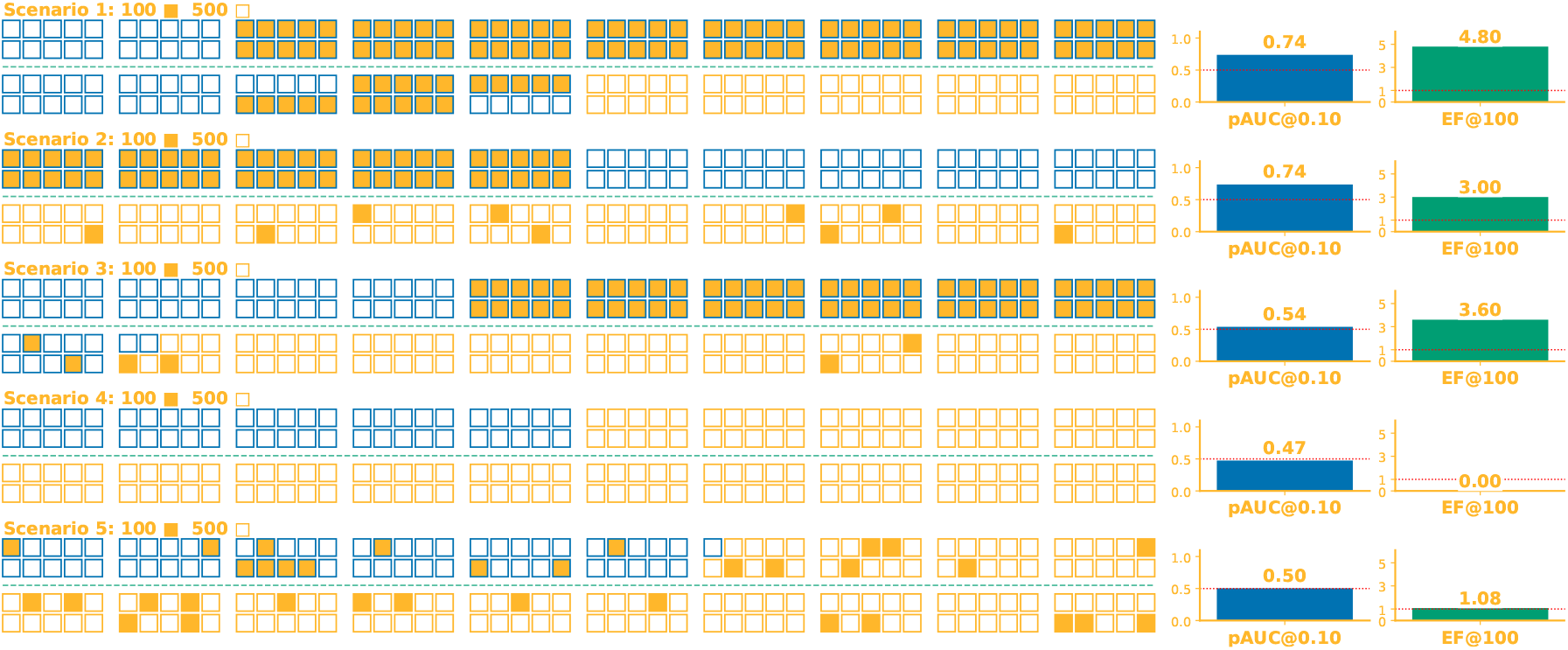
Potential pitfalls of standardized partial AUC (pAUC). Five ranking scenarios compare standardized partial AUC at a false positive rate of 0.10 (pAUC@0.10) with the enrichment factor among the top 100 predictions (EF@100). Each scenario considers for a query TCR, 100 common-specificity TCRs (filled squares) and 500 negative TCRs (empty squares). The horizontal dashed line separates the top 100 ranked TCRs selected for experimental validation; only the first 100 of the remaining 500 candidates are shown below the line. Blue outlines indicate the predictions contributing to pAUC@0.10.

A different evaluation objective arises when TCR similarity methods are used to study the composition of an entire repertoire rather than to rank individual TCRs. In this setting, we are interested in whether TCRs with shared specificity form well-separated clusters. Standard clustering metrics, however, may make assumptions that are not adequate for TCR repertoires. For example, the commonly used Silhouette score evaluates cluster cohesion and separation by comparing the average distance of each TCR to members of the same group with its average distance to the nearest other group [48–50]. When applied to TCR specificity groups, this favours TCRs sharing the same specificity forming a single compact cluster. However, TCRs recognising the same pMHC can contain distinct sequence motifs and form multiple separated sequence motif “islands” [51–54]. In this case, the average distance between TCRs within the same specificity group can be large despite each motif island forming a compact and biologically meaningful local cluster. The Silhouette score may therefore give a poor assessment of specificity clustering when shared specificity TCRs form multiple distinct sequence motif islands.

To address this limitation, we propose a quantile-based local separation score, termed Q-Score, that evaluates local clustering rather than requiring each specificity group to form a single compact cluster. Q-Score measures whether most TCRs within a specificity group have nearby shared specificity TCRs while remaining separated from TCRs with other specificities. The score uses a high quantile of within-specificity nearest-neighbour distances to measure local cohesion and a low quantile of between-specificity distances to measure separation (Supplementary Section S3.5). Using quantiles reduces the influence of individual outliers arising from experimental noise or incorrect annotations, a common source of biases in Silhouette score [55].

Figure 5 illustrates this distinction for TCRs recognising two different epitopes. The GLC-reactive TCRs form two well-separated sequence motif islands, each with distinct CDR3 sequence motifs, despite recognising the same pMHC. Consequently, the Silhouette score is close to zero because the two GLC motif islands are distant from each other, whereas Q-Score remains high because each island forms a compact local cluster separated from TCRs of the other specificity. Q-Score therefore evaluates specificity clustering without assuming that all TCRs recognising the same pMHC belong to a single sequence motif island.

**Figure 5.**
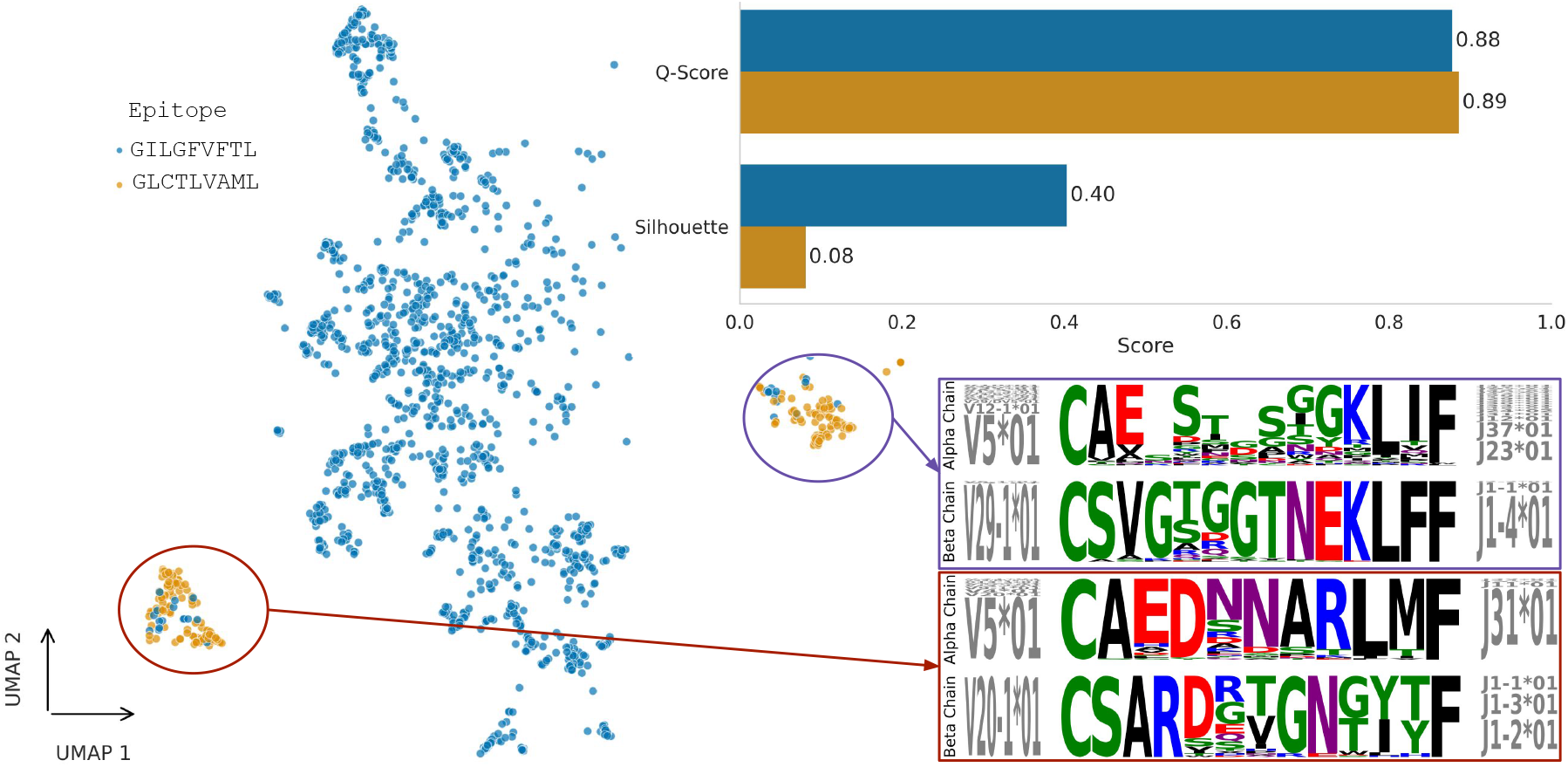
Q-Score evaluates local separation while accommodating multiple sequence motif islands. A UMAP projection of TCR representations for two epitope-specific repertoires illustrates a specificity group that naturally separates into multiple local clusters. Insets highlight representative sequence motif islands, with sequence logos and V/J gene usage shown. The accompanying comparison demonstrates that Q-Score assigns a high score by evaluating local neighborhood purity, whereas the Silhouette score is substantially reduced because it assumes that all TCRs recognizing the same epitope should form a single compact cluster.

AUC, pAUC, EF@*K*, and Q-Score measure different aspects of TCR specificity model performance and should not be treated as interchangeable measures of model quality. AUC is appropriate when overall discrimination across the complete test set is the objective, whereas pAUC focuses on discrimination within a restricted FPR region. When only a limited number of top-ranked candidates are of interest, EF@*K* provides a more direct and intuitive measure of their enrichment, while Q-Score is designed to better accommodate the characteristics of TCR specificity data when evaluating their clustering. Using a metric poorly matched to the intended application may favour the wrong modelling approaches during model development. The choice of evaluation metric should therefore reflect both the biological question being asked and the downstream application of the model.

## 5 Discussion and Conclusion

Despite rapid advances in computational modelling, generalisable TCR–antigen specificity prediction remains an unsolved problem. Our analyses suggest that progress is constrained by the experimental data and the evaluation framework used throughout model development as much as the specific model design. Public repositories of TCR–pMHC data are strongly biased towards a small number of antigens and MHC molecules, sparsely sample the TCR sequence space for individual pMHCs, and contain inconsistencies and errors during data collation. Informative negatives, TCRs that are closely related and experimentally matched but have different specificities, are exceptionally scarce. This leads to the use of uninformative negatives for evaluation, which may result in highly inflated performance. Even when the performance is not inflated, the use of general machine learning evaluation metrics makes interpreting a model’s performance in practical applications difficult and unintuitive. Furthermore, general evaluation metrics such as the Silhouette score may unfairly penalise good TCR models because their underlying assumptions are poorly suited to TCR data and models.

The limitations identified in public TCR–pMHC data cannot be addressed simply by generating more records. New data should instead be generated to address specific gaps in the available TCR–pMHC data, including poorly represented antigens and HLAs, and limited sampling of TCR diversity for individual pMHCs. The TCR–pMHC data should contain complete paired-chain TCR information and, where possible, be paired with informative negatives.

Improving the available data alone is insufficient if model evaluation does not measure the biological properties we want models to capture. The choice of evaluation metric therefore requires careful consideration to ensure that it is intuitive, suitable for TCR data, and aligned with the downstream application of the model. These choices are particularly consequential because model evaluation guides future model development by determining which approaches appear most promising and are prioritised for further improvement. A misleading evaluation can therefore favour modelling approaches that perform well for the wrong reasons or according to criteria that do not match the intended downstream application. The data and the evaluation framework are therefore crucial for determining when models can be trusted, understanding their limitations, and directing future model development towards generalisable TCR–pMHC specificity prediction.

Addressing these challenges will require coordinated community effort across experimental data generation, database curation, benchmark construction, and evaluation metrics. The success of protein structure prediction provides an example of what such sustained community efforts can enable. Large quantities of consistently represented experimental data in the Protein Data Bank (PDB), combined with robust and fair benchmarking through CASP, provided the necessary foundations for the development of breakthroughs such as AlphaFold [56–58]. Recent studies reflect growing recognition of these challenges and the beginnings of such efforts within the TCR community. Messemaker et al. [37] and Gfeller et al. [26], among others, highlight the problems with the data, while Minervina et al. [27] and Lu et al. [22] provide an indication of the importance of informative negatives and paired-chain data. At the same time, Gaglione et al. [59] introduce a scalable experimental approach for TCR synthesis and antigen screening. The approach provides a practical route towards generating larger, more representative datasets with high confidence and informative negatives generated during the screening process. As novel technologies generate increasingly large and heterogeneous datasets, assay harmonisation and standardisation are essential for generating reliable and comparable data across studies [60, 61]. Such foundations would enable models to be trained on higher quality and more representative data, while providing the robust benchmarks needed to identify genuine improvements in generalisability. Ultimately, progress towards generalisable TCR–pMHC specificity prediction requires looking beyond better models and improving the data and evaluation foundations on which their development depends.

## Author Contributions

**I.I**. and **H.K**. conceptualized the study. **I.I**. developed the methodology and software, curated the data, performed the formal computational analyses, generated the visualizations, and wrote the original draft of the manuscript. **A.A**. and **G.O**. provided immunological insights, evaluated the clinical relevance of the benchmarks, and contributed to the investigation. **H.K**. supervised the project and acquired funding. All authors critically reviewed, edited, and approved the final manuscript.

## Acknowledgements

This work was supported by funding from the UK Medical Research Council grant number MC UU 12010/3 to I.I. and H.K. A.A. is supported by the Wellcome Trust (Grant No. 315684/Z/24/Z). G.O. is supported by the Medical Research Council.

## Data and code availability

All of the code and data used to conduct this analysis can be found at: https://github.com/ili3p/generalisable-tcr-antigen-prediction.

