## Supplementary document for "Beyond Better Models: Barriers and a Path Forward in Generalisable TCR–Antigen Specificity Prediction"

#### Supplementary Material

Ilija Ilievski[1,2], Agne Antanaviciute[1,2], Graham Ogg[1], and Hashem Koohy[1,2,†]

[1]Medical Research Council Translational Immune Discovery Unit (MRC TIDU), Weatherall Institute of Molecular Medicine, University of Oxford, United Kingdom
[2]MRC WIMM Centre for Computational Biology, Weatherall Institute of Molecular Medicine, Oxford, United Kingdom

#### S1 Experimental Setup

We evaluated SCEPTR, available at `https://sceptr.readthedocs.io/`, a contrastively pretrained TCR representation model [1], and TCRdist3, Python 3 implementation of TCRdist [2, 3], available at `https://tcrdist3.readthedocs.io/`.

SCEPTR was used without task-specific fine-tuning. Distances between groups of TCRs were obtained with `sceptr.calc_cdist_matrix`.

TCRdist3 was used with its standard TCR distance calculation.

Thus, neither method was trained or calibrated on the Croce et al. binding labels before evaluation [4].

#### S2 Source code

The code for the group-based evaluation described in Section 2, preprocessing of the FASTQ files from Croce et al., IEDB and VDJDb data analysis, and calculation of the evaluation metrics is available at `https://github.com/ili3p/generalisable-tcr-antigen-prediction`.

### S3 Evaluation Metric Definitions

#### S3.1 Area Under the Receiver Operating Characteristic Curve (AUC) calculation

When applied to evaluating similarity (distance) models, the AUC score is calculated as the probability that a randomly selected positive distance is smaller than a randomly selected negative distance. For example, in Section 3, the positive distances for group 1 and 2 are the distances between each unique pair of TCRs from the group of binders. The negative distances for group 1 and 2 are the distances between each unique pair of TCRs where one TCR is a binder and the other is non-binder. The positive distances for group 3 are the distances between each unique pair of TCRs where one TCR is a mutational scanning binder from Croce et al. and the other is a VDJdb binder excluding those from Croce et al. The negative distances for group 3 are the distances between each unique pair of TCRs where one TCR is a mutational scanning binder from Croce et al. and the other is a mutational scanning non-binder from Croce et al.

Specifically, for a distance threshold $\tau$, the true positive rate (TPR) represents the probability that a positive distance $d^+$ is smaller than or equal to $\tau$. Similarly, the false positive rate (FPR) represents the probability that a negative distance $d^-$ is smaller than or equal to $\tau$. These quantities are defined as

$$\mathrm{TPR}(\tau) = \Pr\bigl(d^+ \leq \tau\bigr), \qquad \mathrm{FPR}(\tau) = \Pr\bigl(d^- \leq \tau\bigr).$$

The ROC curve is then obtained by varying $\tau$, and the area under this curve is given by

$$\mathrm{AUC} = \int_0^1 \mathrm{TPR}\bigl(\mathrm{FPR}^{-1}(u)\bigr)\, \mathrm{d}u = \Pr\bigl(d^+ < d^-\bigr),$$

Thus, the AUC can be interpreted as the probability that a randomly selected positive pair has a smaller distance than a randomly selected negative pair [5].

#### S3.2 Partial AUC

Partial AUC (pAUC) restricts the evaluation to a selected region of the ROC curve, typically the region with a false positive rate below a predefined threshold $\alpha$:

$$\mathrm{pAUC}_\alpha = \int_0^\alpha \mathrm{TPR}\bigl(\mathrm{FPR}^{-1}(u)\bigr)\, \mathrm{d}u.$$

Thus, $\mathrm{pAUC}_\alpha$ measures the discriminative performance over the portion of the ROC curve corresponding to $\mathrm{FPR} \leq \alpha$.

Partial AUC must be standardised to map the raw values to the familiar scale on which 0.5 represents random ranking and 1 represents perfect ranking. A commonly used standardisation is the McClish standardization [6, 7].

$$\mathrm{pAUC}_\alpha^{\mathrm{std}} = \frac{1}{2}\left[1 + \frac{\mathrm{pAUC}_\alpha - \frac{1}{2}\alpha^2}{\alpha - \frac{1}{2}\alpha^2}\right]. \tag{1}$$

Although this transformation maps random performance to 0.5 and perfect performance to 1, it does not map a completely reversed ranking to 0. For example, when $\alpha = 0.1$, a perfectly reversed ranking receives a standardised partial AUC of approximately 0.474. Thus, values close to 0.5 do not necessarily imply random performance after standardisation.

#### S3.3 Enrichment Factor at K

The enrichment factor at rank $K$ is defined as:

$$\text{EF@}K = \frac{\frac{m}{K}}{\frac{n}{N}} = \frac{mN}{nK}, \tag{2}$$

where $N$ is the total number of TCRs (shared-specificity TCRs plus negative TCRs), $n$ is the number of shared-specificity TCRs within this set, $K$ is the number of top-ranked TCRs considered, and $m$ is the number of shared-specificity TCRs among these $K$ TCRs.

The numerator, $m/K$, is the observed fraction of shared-specificity TCRs among the top $K$ TCRs, whereas $n/N$ is the fraction expected from the complete TCR set. An enrichment factor of 1 therefore corresponds to random retrieval, where the proportion of shared-specificity TCRs among top $K$ is equal to that expected given their prevalence in the complete TCR set. An enrichment factor greater than 1 indicates enrichment in the top $K$, and values below 1 indicate depletion.

#### S3.4 Silhouette Score

The Silhouette score measures how well each TCR is separated from TCRs belonging to other groups relative to how closely it is grouped with TCRs from the same group [8]. For a TCR $i$, let $a(i)$ denote the mean distance between $i$ and all other TCRs belonging to the same group:

$$a(i) = \frac{1}{|C_i| - 1} \sum_{\substack{j \in C_i \\ j \neq i}} d(i, j), \tag{3}$$

where $C_i$ denotes the group containing TCR $i$, $|C_i|$ is the number of TCRs in that group, and $d(i, j)$ is the distance between TCRs $i$ and $j$.

For every other group $C \neq C_i$, the mean distance between TCR $i$ and the TCRs in $C$ is calculated. The smallest of these mean distances is denoted by $b(i)$:

$$b(i) = \min_{C \neq C_i} \left[ \frac{1}{|C|} \sum_{j \in C} d(i, j) \right]. \tag{4}$$

Thus, $a(i)$ measures the average within-group distance, whereas $b(i)$ measures the average distance to the nearest other group.

The Silhouette score for TCR $i$ is then defined as

$$s(i) = \frac{b(i) - a(i)}{\max\{a(i), b(i)\}}, \tag{5}$$

with $s(i) \in [-1, 1]$. A value close to 1 indicates that the TCR is substantially closer to TCRs in its own group than to those in the nearest other group. A value close to 0 indicates that the TCR lies approximately equally close to its own group and the nearest other group, whereas a negative value indicates that it is, on average, closer to another group than to its assigned group.

The overall Silhouette score is obtained by averaging the individual Silhouette scores across all $N$ TCRs:

$$\text{Silhouette} = \frac{1}{N} \sum_{i=1}^{N} s(i). \tag{6}$$

##### S3.5 Q-Score

To evaluate local compactness without requiring each specificity group to form a single global cluster, we propose a robust, non-parametric, quantile-based local separation score, termed Q-Score.

For each TCR $i$ belonging to specificity group $g$, let

$$d_{i,(k)}^{\text{intra}} \tag{7}$$

denote the distance from $i$ to its $k$-th nearest TCR with shared specificity. The group-level intra-specificity distance is defined as the 0.95-quantile of these distances:

$$D_g^{\text{intra}}(k) = Q_{0.95}\left(\left\{d_{i,(k)}^{\text{intra}} \,:\, i \in g\right\}\right), \tag{8}$$

where $Q_q$ denotes the $q$-quantile. This quantity provides a robust measure of local compactness within specificity group $g$, with smaller values indicating that most TCRs have close shared-specificity neighbours.

To quantify separation from other specificity groups, we consider all pairwise distances between TCRs belonging to group $g$ and TCRs belonging to any other specificity group:

$$D_g^{\text{inter}} = Q_{0.05}\left(\{d(i,j) \,:\, i \in g,\ j \notin g\}\right). \tag{9}$$

Here $Q_{0.05}$ characterises the lower tail of the inter-specificity distances and therefore makes the score sensitive to regions in which TCRs from different specificity groups approach or overlap one another, while reducing sensitivity to extreme distances.

The relative local compactness and separation of specificity group $g$ is then quantified by

$$R_g(k) = \frac{D_g^{\text{intra}}(k)}{D_g^{\text{inter}}} = \frac{Q_{0.95}\left(\left\{d_{i,(k)}^{\text{intra}} \,:\, i \in g\right\}\right)}{Q_{0.05}\left(\{d(i,j) \,:\, i \in g,\ j \notin g\}\right)}. \tag{10}$$

Smaller values of $R_g(k)$ indicate better local compactness and separation, as the within-specificity neighbourhood distances are small relative to the lower tail of the between-specificity distances.

For a high-is-better representation, the ratio is transformed monotonically as

$$S_g(k) = \frac{1}{1 + R_g(k)}, \tag{11}$$

such that $S_g(k) \in [0,1]$, with values approaching 1 indicating strong local compactness and separation, and values approaching 0 indicating poor separation. The overall Q-Score is defined as the mean of the group-level scores across all $G_{\text{valid}}$ specificity groups that contain at least $k+1$ TCRs:

$$\text{Q-Score}(k) = \frac{1}{|G_{\text{valid}}|} \sum_{g \in G_{\text{valid}}} S_g(k). \tag{12}$$

Specificity groups containing fewer than $k+1$ TCRs are excluded because the $k$-th nearest shared-specificity neighbour is undefined. In all our experiments we set $k = 10$.

#### S4 Supplementary Figures

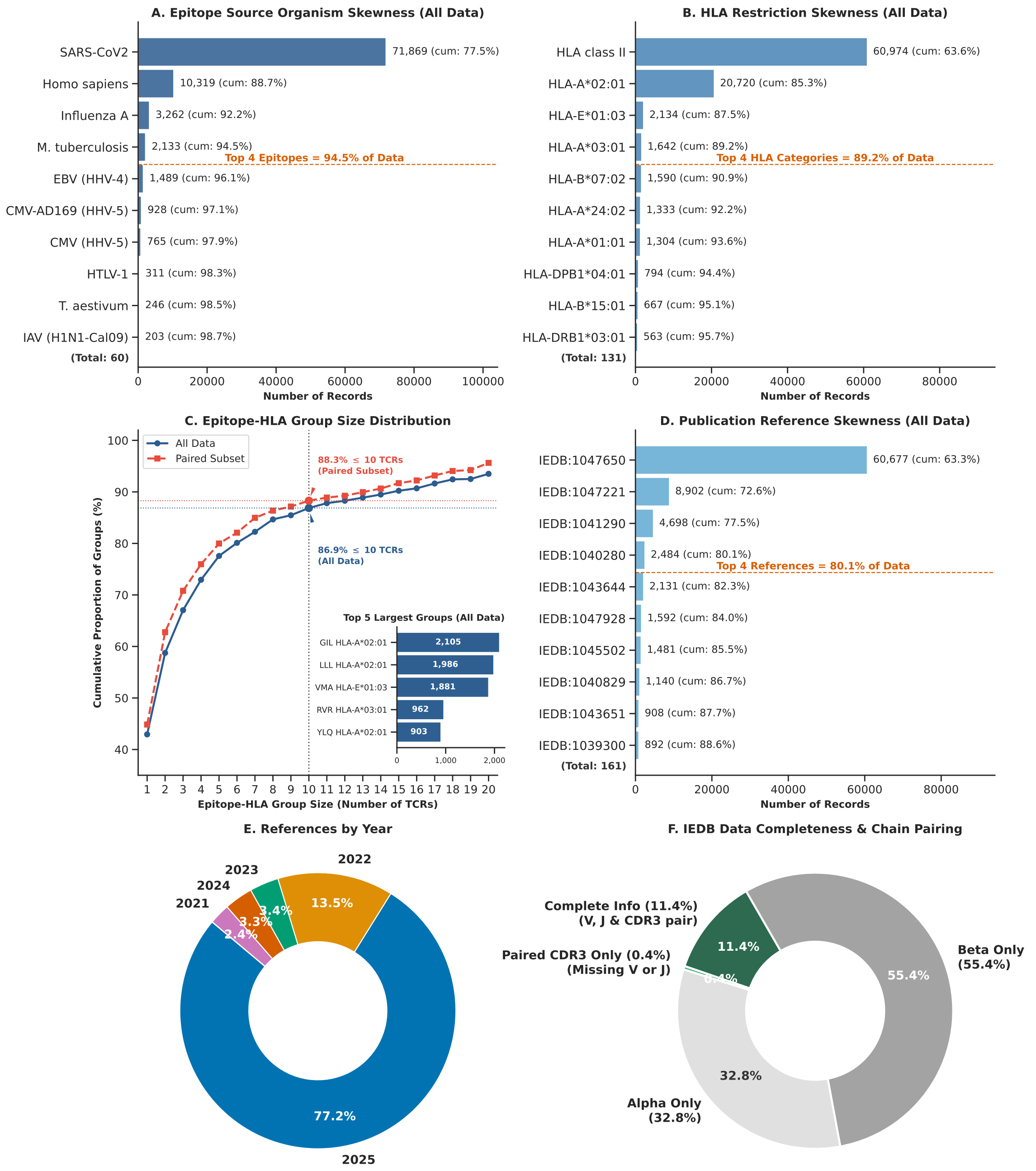


Figure S1: **Quantitative analysis of data biases and annotation incompleteness in human αβ TCR–pMHC data from IEDB published in 2021 or later (≈95K records)** (**A**) Epitope skewness. (**B**) HLA restriction skewness. (**C**) Cumulative distribution of number of unique TCR sequences per epitope–HLA pair. (**D**) Publication skewness. (**E**) Publication year distribution. (**F**) Data completeness and chain pairing.

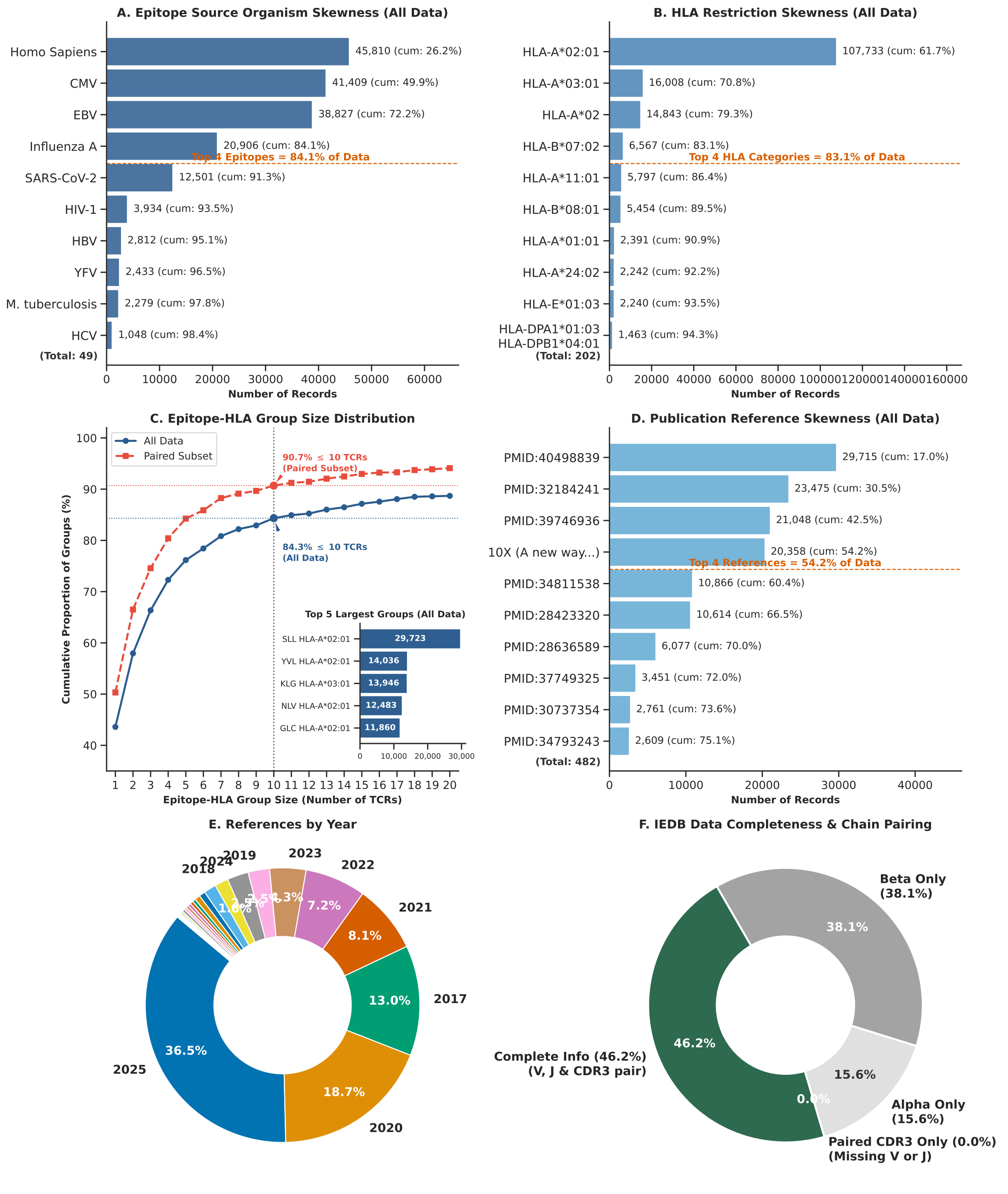


Figure S2: **Quantitative analysis of data biases and annotation incompleteness in human αβ TCR–pMHC data from VDJdb (≈174K records)** (**A**) Epitope skewness. (**B**) HLA restriction skewness. (**C**) Cumulative distribution of number of unique TCR sequences per epitope–HLA pair. (**D**) Publication skewness. (**E**) Publication year distribution. (**F**) Data completeness and chain pairing.
